# TGF-β signaling regulates epithelial permeability in Drosophila ovaries by modulating adhesion independent of actomyosin contractility

**DOI:** 10.64898/2026.02.14.705893

**Authors:** Harshath Amal, Thea Jacobs, Max Lohrberg, Stefan Luschnig

## Abstract

Epithelial morphogenesis and homeostasis rely on dynamic remodeling of cell-cell junctions. Tricellular junctions (TCJs) at cell vertices are key sites that control epithelial permeability and plasticity, yet how TCJs are remodeled remains unclear. In the follicular epithelium (FE) in Drosophila ovaries, TCJs open transiently in a process called patency to allow passage of yolk proteins for uptake by the oocyte. We investigated how a gradient of TGF-β signaling activity suppresses patency in a graded manner across the FE. We show that TGF-β signaling blocks patency in a cell-autonomous manner by strengthening E-Cadherin (E-Cad)-based adhesion through inducing E-Cad transcription and preventing its removal from cell vertices. In parallel, TGF-β signaling activates myosin II through Rho-Rok signaling. However, myosin II activity is dispensable for TGF-β-mediated suppression of patency. We show that TGF-β signaling controls TCJ remodeling in follicle cells primarily by reinforcing E-Cad-based adhesion, in part through upregulating p120-catenin. Our findings disentangle the roles of adhesion and actomyosin contractility in maintaining TCJ integrity and reveal how a tissue-scale morphogen gradient spatially patterns epithelial permeability during development.

## Introduction

Normal organ development and function rely on the dynamic remodeling of intercellular contacts, enabling tissues to reshape during homeostasis and in response to infection, inflammation, and injury (Mira-Osuna and Le Borgne, 2024). In epithelial and endothelial barriers, junctional permeability is regulated to control paracellular transport of solutes and transmigration of immune cells. Conversely, aberrant weakening of cell-cell junctions is a hallmark of infections, inflammatory conditions, and cancer metastasis, highlighting the critical role of junction integrity in preserving barrier function and tissue homeostasis.

While bicellular junctions (BCJs) connecting two cells are the most abundant intercellular contacts, tricellular junctions (TCJs) at cell vertices have emerged as key sites that control epithelial permeability and plasticity (Bosveld et al., 2018; Higashi and Miller, 2017). TCJs play vital roles in cell shape, cytoskeletal organization, spindle orientation, and epithelial barrier function. They comprise a distinct set of adhesion proteins and are dynamically remodeled during cell rearrangements, immune cell transmigration, and infection processes (Furuse et al., 2014; Higashi and Chiba, 2020; Higashi and Miller, 2017). However, how epithelial cells modulate TCJ permeability without compromising tissue integrity is not well understood.

Ovarian development in Drosophila provides an accessible model to address these questions. The ovary consists of chains of follicles, each comprising 16 germline cells (15 nurse cells and one oocyte) surrounded by a single-layered somatic follicular epithelium (FE; Berg et al., 2023; Horne-Badovinac and Bilder, 2005). During vitellogenesis (yolk uptake) in mid-oogenesis, cell vertices in the FE open transiently to generate intercellular channels that allow passage of hemolymph-borne yolk proteins to reach the oocyte surface for endocytic uptake. This process, known as follicular patency (Patchin and Davey, 1968; Pratt and Davey, 1972), is conserved among oviparous animals and offers a tractable system for dissecting the mechanisms of TCJ remodeling. Patency is initiated by endocytosis-dependent removal of adhesion proteins (E-Cad, N-Cad, NCAM/Fasciclin 2 (Fas2), Sidekick (Sdk)) from follicle cell (FC) vertices and downregulation of actomyosin contractility, and is terminated by the assembly of occluding septate junctions that seal the gaps between FCs (Isasti Sanchez et al., 2021; Jacobs et al., 2025).

A key question is which signals and cellular mechanisms control TCJ remodeling during patency. The local activation of signaling pathways (BMP, EGFR, JAK/STAT) subdivides the FE into distinct regions and cell fates, including anterior (stretched) follicle cells covering the nurse cells and different types of main body follicle cells (MBFCs) overlying the oocyte (Fig. 1A; Berg et al., 2023; Duhart et al., 2017). Activation of BMP and EGFR signaling in the dorsal-anterior region of the main body follicular epithelium specifies a subset of FCs referred to as centripetal FCs (CPFCs) that express elevated levels of adhesion proteins (E-Cad, Fasciclin 3; Niewiadomska et al., 1999; Shravage et al., 2007) and give rise to anterior structures of the mature eggshell (Osterfield et al., 2013). Unlike the rest of the MBFCs, the CPFCs display little or no opening of vertices during patency (Row et al., 2021). Moreover, ectopic activation of BMP signaling throughout the FE was shown to block patency, whereas reduced EGFR activation caused ectopic patency in the dorsal-anterior region, suggesting that BMP and EGFR signaling regulate processes that control vertex opening (Row et al., 2021).

**Figure 1.**
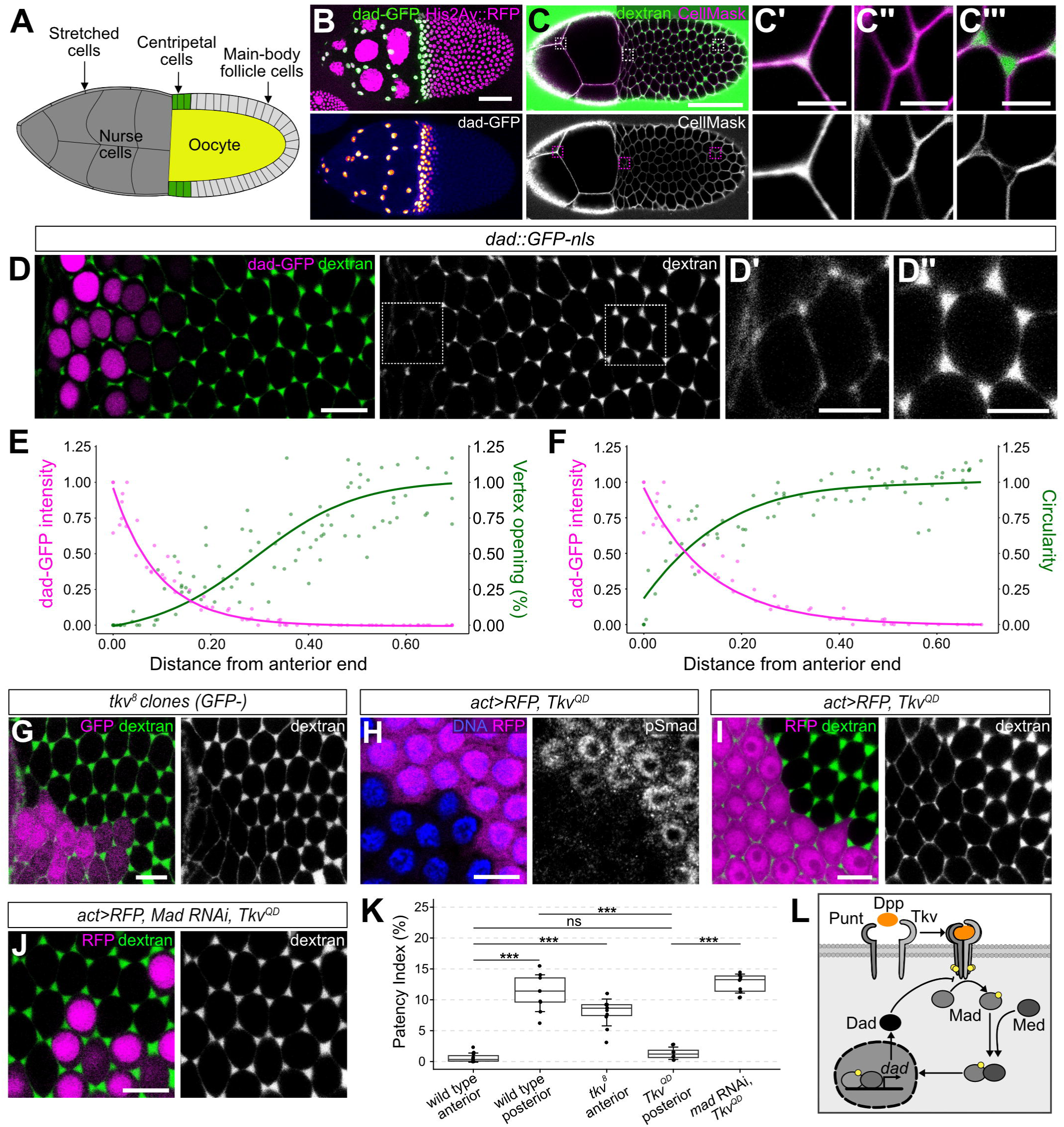
Graded TGF-β signaling regulates vertex opening across the follicular epithelium. (A) Scheme of patent (stage 10A) follicle showing cell types referred to in the text. Anterior is to the left. (B) Confocal Projection of fixed follicle expressing *dad*::GFP-nls (green) and His2Av::RFP (magenta). Single channel shows *dad*::GFP-nls gradient with intensities color-coded using heatmap. Specimens in this and all subsequent panels are in mid-patency (stage 10A), unless indicated otherwise. (C) Confocal section at basolateral level in living follicle. Dextran (green) marks intercellular gaps, CellMask (magenta) marks plasma membranes. Note that intercellular gaps are absent between stretched cells (C’) and centripetal cells (C’’) but present between posterior MBFCs (C’’’). (D) Basolateral section of living follicle expressing *dad*::GFP-nls (magenta). Note that dextran-filled intercellular gaps (green) are absent or small between anterior MBFCs (*dad*::GFP-nls–positive nuclei) at the nurse cell–MBFC boundary (left) but larger in posterior FE (right). Insets highlight vertices in anterior (D’) and posterior (D’’) regions. (E) Quantification of *dad*::GFP-nls signal (magenta curve) and vertex opening size (green curve) across the main body FE as a function of distance from the nurse cell–MBFC boundary. (F) Quantification of *dad*::GFP-nls signal (magenta curve) and circularity of MBFC cross-section (green curve) across the main body FE as a function of distance from the nurse cell–MBFC boundary. (G) Section of living follicle carrying *tkv^8^* clone marked by absence of GFP (magenta). Dextran (green) marks intercellular gaps. Note larger gaps in *tkv^8^* clone compared to control cells. (H) Section of fixed follicle expressing Tkv^QD^ in RFP-marked cells (magenta). Nuclei are labelled with Hoechst (blue). Activated SMAD (pSMAD; single-channel image) indicates active TGF-β signaling in Tkv^QD^-expressing cells. (I) Section of living follicle expressing Tkv^QD^ in RFP-marked cells (magenta). Note smaller intercellular gaps (dextran, green) between Tkv^QD^-expressing cells compared to control cells. (J) Section of living follicle expressing Tkv^QD^ and *mad* dsRNA in RFP-marked cells (magenta). Note restoration of normal intercellular gap size by *mad* RNAi in Tkv^QD^-expressing cells. (K) Quantification of intercellular gap size (patency index) in anterior or posterior FE regions in the indicated genotypes. p-values (Pairwise Wilcoxon Rank Sum Test): *** p ≤ 0.001; ns, not significant. (L) Scheme of TGF-β signaling pathway in *Drosophila melanogaster*. Scale bars: (B,C), 50 µm; (C’-C’’’), 5 µm; (D) 10 µm; (D’, D’’), 5 µm; (G-J), 10 µm.

To understand how TCJ permeability is regulated, we investigated how BMP signaling acts in FCs to block patency. The BMP ligand Decapentaplegic (Dpp) activates canonical BMP signaling through the type I receptors Thickveins (Tkv) and Saxophone (Sax) and the type II receptor Punt (Put). Receptor activation leads to phosphorylation of the intracellular signal transducer and transcription factor Mothers against Dpp (Mad), which then translocates with its binding partner Medea to the nucleus and regulates transcription of target genes, mainly through repressing the BMP antagonist Brinker (Akiyama et al., 2023). This pathway is essential for maintenance and proliferation of germline stem cells in the germarium (Xie and Spradling, 1998), and later patterns the FE during mid-oogenesis. Dpp secreted by the anterior stretched FCs forms an anterior-to-posterior gradient in the oocyte-associated FCs and determines distinct FC fates, in part through cross-talk with EGFR signaling (Charbonnier et al., 2015; Chen and Schüpbach, 2006; Osterfield et al., 2013; Shravage et al., 2007; Yakoby et al., 2008). Dpp signaling was shown to promote remodeling of adherens junctions (AJs) and relocalization of Myosin II, enabling flattening of stretched FCs (Brigaud et al., 2015) and collective migration of CPFCs (Niewiadomska et al., 1999; Parsons et al., 2023). However, how Dpp signaling regulates TCJ integrity was not known.

Here we show that BMP signaling acts in a cell-autonomous manner to prevent vertex opening by inducing actomyosin contractility through Rho signaling, upregulation of E-Cad transcription, and retention of E-Cad at FC vertices. We found that while E-Cad-based adhesion is crucial for the TGF-β-mediated suppression of patency, Rho signaling and MyoII activity are not required. TGF-β signaling acts primarily by reinforcing E-Cad-based adhesion, in part through upregulating p120-catenin. Our findings reveal how a morphogen controls epithelial permeability through regulating vertex-specific cell adhesion.

## Results

To analyze the spatial pattern of TGF-β signaling activity across the FE, we used a transcriptional reporter expressing nuclear-localized GFP under the control of TGF-β-responsive regulatory sequences of the *daughters against dpp* (*dad*) gene (*dad*::GFP-nls; Ninov et al., 2010). Nuclei of the stretched FCs covering the nurse cells showed the highest *dad*::GFP-nls signals in patent follicles (stage 10A; Fig. 1B; Charbonnier et al., 2015). Additionally, the CPFCs overlying the anterior margin of the oocyte displayed *dad*::GFP-nls signals, which declined in a steep anterior-to-posterior gradient (Fig. 1B,D, Fig. 2A,B). This gradient was inversely correlated with the size of intercellular openings, which were small or undetectable in the stretched FCs and in CPFCs, and increased in size towards the posterior to reach maximum size in the FE covering the oocyte (Fig. 1C,D,E). Along the same anterior-posterior gradient, the outlines of FCs changed from hexagonal to rounded, as quantified by the circularity of the FC cross-sectional area (Fig. 1D,F). Consistent with these findings, CPFCs resisted the transient opening of vertices induced by hyperosmotic treatment (Jacobs et al., 2025), whereas more posterior MBFCs readily opened their vertices upon hyperosmotic treatment with DMSO (MovieLS1).

**Figure 2.**
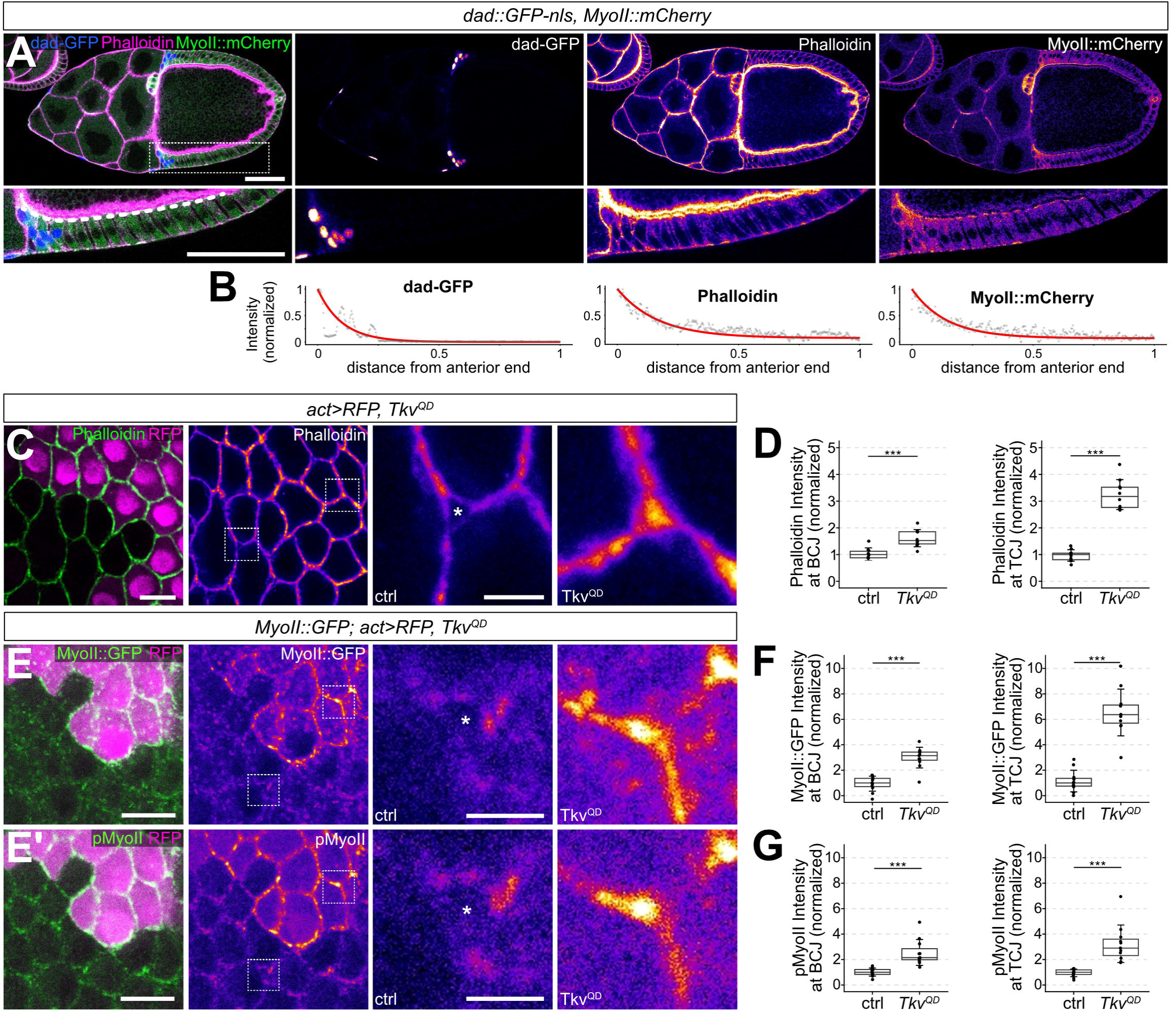
TGF-β signaling regulates actomyosin contractility in follicle cells. (A) Mid-sagittal section of fixed follicle expressing *dad*::GFP-nls (magenta) and MyoII::mCherry (green), stained with Phalloidin (F-actin; magenta). Intensities are color-coded using heatmap in single-channel images. Lower panels show closeup of region marked by box in upper left panel. (B) Normalized fluorescence intensities measured along line profile (dashed line in (A)). (C) Basolateral section of fixed follicle expressing Tkv^QD^ in RFP-marked cells (magenta). Phalloidin labels F-actin (green in merge, heatmap in single-channel images). Closeups of boxed vertices show elevated levels of F-actin at bicellular junctions (BCJ) and tricellular junctions (TCJ) in Tkv^QD^-expressing cells compared to control (ctrl) cells. (D) Quantification of Phalloidin signals at BCJs and TCJs in control and Tkv^QD^-expressing clones. p-values (Welch’s two-sample t-test): *** p ≤ 0.001. (E,E’) Basolateral section of fixed follicle expressing MyoII::GFP and Tkv^QD^ in RFP-marked cells (magenta). Single-channel images show MyoII::GFP (E) or pMyoII staining (E’). Closeups show single vertices of wild-type (ctrl) or Tkv^QD^-expressing cells. Note higher levels and vertex accumulation of MyoII (E) and pMyoII (E’) and vertex accumulation in Tkv^QD^-expressing cells. (F) Quantification of MyoII signals at BCJs or TCJs in control and Tkv^QD^ expressing clones. P-values (Welch’s two-sample t-test): *** p ≤ 0.001. (G) Quantification of pMyoII signals at BCJs or TCJs in control and Tkv^QD^ expressing clones. P-values (Welch’s two-sample t-test): *** p ≤ 0.001. Scale bars: (A), 50 µm; (C, E, E’), 10 µm, closeups 2.5 µm. Asterisks mark open vertices in (C,E,E’).

### TGF-**β** signaling prevents remodeling of tricellular junctions during patency

These findings suggest that TGF-β signaling suppresses vertex remodeling and cell shape changes in the anterior FE during patency. Consistent with this notion, loss of the type I TGF-β receptor Thickveins (Tkv) in *tkv^8^* mutant clones located in the CPFC region led to ectopic vertex opening in the Tkv-deficient cells (Fig. 1G,K). Conversely, expression of a constitutively activated version of the Tkv receptor (Tkv^Q253D^, referred to as Tkv^QD^; Nellen et al., 1996) in FC clones located in the posterior FE led to accumulation of phosphorylated SMAD (pSMAD) in the nucleus (Fig. 1H), indicative of active TGF-β signaling, and prevented vertex opening in a cell-autonomous fashion (Fig. 1I,K). The block of patency in Tkv^QD^-expressing cells was suppressed by RNAi-induced depletion of the intracellular TGF-β signal transducer Mad (Fig 1J,K). Together, these findings indicate that activation of signaling through the canonical TGF-β signaling pathway (Fig. 1L) is necessary and sufficient to suppress vertex opening in the FE.

### TGF-**β** signaling regulates actomyosin contractility via Rho-Rok signaling

To understand how TGF-β signaling controls vertex opening, we investigated its effects on cell-cell adhesion and actomyosin contractility. Anterior FCs (stretched FCs and centripetal cells) showed elevated levels of cortical F-actin and non-muscle myosin II (MyoII), with intensities mirroring the anterior-posterior gradient of *dad*::GFP-nls expression (Fig. 2A,B). Similarly, the levels of E-Cad were elevated at AJs of anterior FCs and decreased in a gradient towards the posterior end (Fig. 4C). These findings suggest that TGF-β signaling might control vertex opening through regulating actomyosin contractility and cell-cell adhesion in FCs.

To test whether activation of TGF-β signaling can induce actomyosin contractility, we analyzed the distribution and levels of F-actin and MyoII in Tkv^QD^-expressing FC clones. Compared to neighboring control cells, Tkv^QD^-expressing cells showed significantly elevated levels of cortical F-actin (Fig. 2C,D) and of GFP-tagged MyoII (MyoII::GFP) at BCJs and, more strongly, at TCJs (Fig. 2E,F). Staining with anti–phospho-MyoII (pMyoII) antibodies, which detect the phosphorylated myosin regulatory light chain (MRLC), revealed a corresponding increase in active MyoII (Fig. 2E’,G). The ratio of active MyoII (pMyoII) to total MyoII (MyoII::GFP) at BCJs and TCJs remained unchanged between Tkv^QD^-expressing cells and control cells, suggesting that TGF-β signaling promotes actomyosin contractility by both upregulating MyoII levels and enhancing MyoII activity through MRLC phosphorylation. Consistent with these results, Tkv^QD^-expressing cells showed elevated levels of Rho1, a key regulator of MyoII activity (Fig. 3A,B).

**Figure 3.**
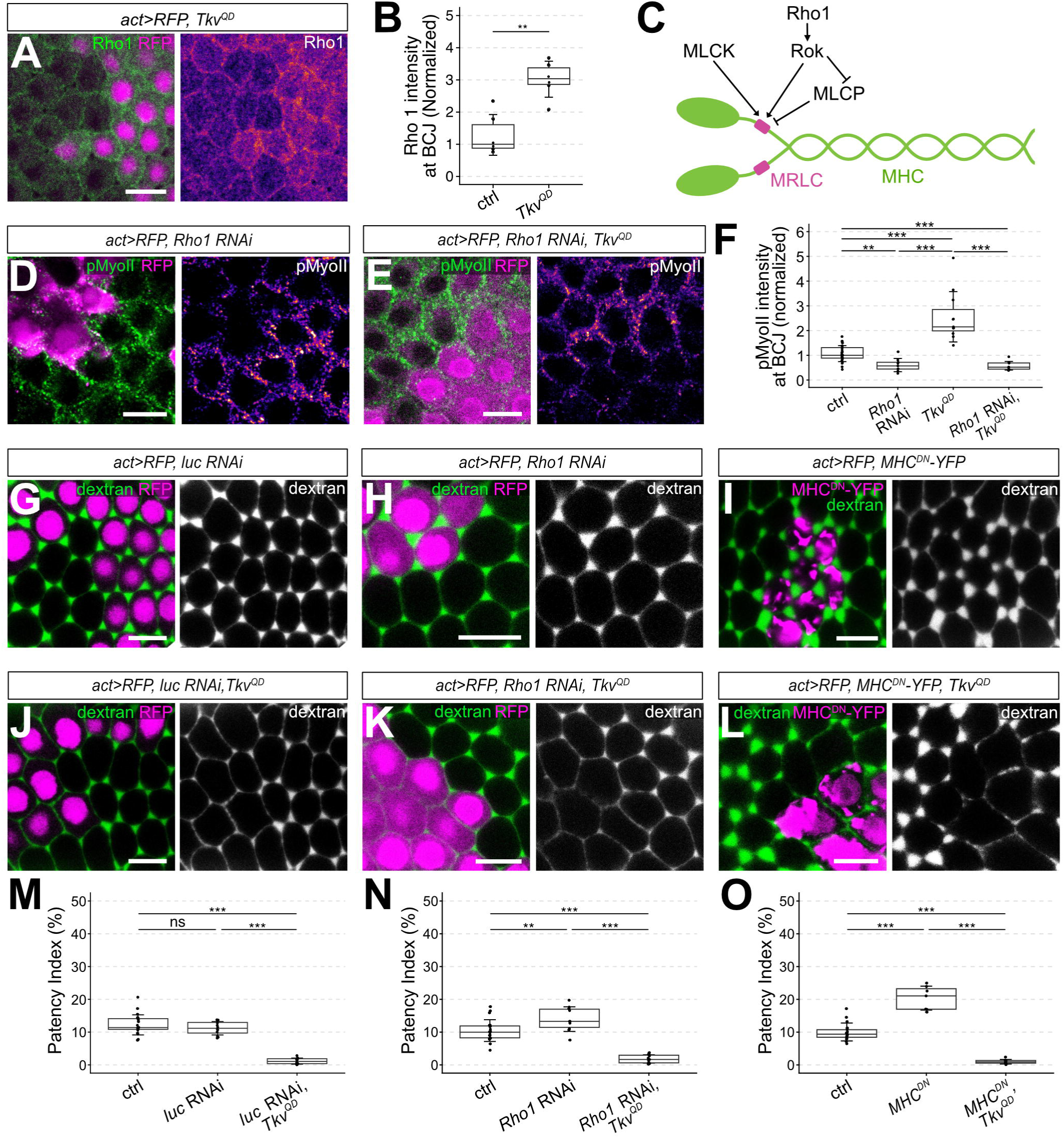
Myosin II activity is dispensable for TGF-β mediated suppression of patency. (A) Basolateral section of fixed follicle expressing Tkv^QD^ in RFP-marked cells (magenta) stained for Rho1. (B) Quantification of Rho1 signals at BCJs. p-values (Welch’s two-sample t-test): ** p ≤ 0.01. (C) Scheme showing regulation of myosin II activity through phosphorylation of myosin regulatory light chain (MRLC). MHC, myosin heavy chain; MLCK, myosin light-chain kinase; MLCP, myosin light-chain phosphatase; Rok, Rho-associated kinase. (D,E) Basolateral sections of follicles expressing *Rho1* dsRNA (D) or Tkv^QD^ and *Rho1* dsRNA (E) in RFP-marked cells (magenta). Anti-pMyoII staining (green) detects active MyoII. (F) Quantification of pMyoII signals at BCJs in the indicated genotypes. p-values (Pairwise Wilcoxon Rank Sum Test): ** p ≤ 0.01; *** p ≤ 0.001. (G-I) Basolateral sections of living follicles expressing *luciferase* (*luc;* control) dsRNA (G), *Rho1* dsRNA (H) or MHC^DN^-YFP (I) in RFP-marked cells (magenta). Dextran (green) marks intercellular gaps. Note enlarged gaps in Rho1-depeted cells (H) and MHC^DN^-YFP-expressing cells (I). (J-L) Basolateral sections of living follicles expressing Tkv^QD^ along with *luciferase* dsRNA (control; J), *Rho1* dsRNA (K) or MHC^DN^-YFP (L) in RFP-marked cells (magenta). Note that neither depletion of Rho1 (K) nor inactivation of MyoII (L) restore vertex opening in Tkv^QD^-expressing cells. (M-O) Quantification of patency index in the indicated genotypes. p-values (Pairwise Wilcoxon Rank Sum Test): ** p ≤ 0.01; *** p ≤ 0.001; ns, not significant. Scale bars: (A, D, E, G-L), 10 µm. See also Figure S1.

### TGF-**β**-induced MyoII activity is not required to prevent vertex opening

Previous work demonstrated that reduction of actomyosin contractility at FC vertices is necessary for patency initiation (Isasti Sanchez et al., 2021), whereas excessive MyoII activity upon overactivation of Rho signaling inhibits vertex opening (Jacobs et al., 2025). Hence, our findings suggest that TGF-β signaling blocks patency by upregulating MyoII levels and activity to induce actomyosin contractility. We therefore asked whether suppressing MyoII activity could restore vertex opening in the presence of activated TGF-β signaling. To suppress MyoII activation, we depleted Rho1 or its downstream effector Rho-associated kinase (Rok; Fig. 3C; Amano et al., 2010) by RNAi in FC clones. This led to substantial reduction of active MyoII (pMyoII; Fig. 3D,F, Fig. S1) and a corresponding increase in intercellular gap size within the RNAi clones (Fig. 3H,N, Fig. S1), whereas a control RNAi construct (luciferase RNAi) had no detectable effect (Fig. 3G,M). In Tkv^QD^-expressing FC clones, depleting Rho1 or Rok reduced pMyoII to levels comparable to those upon Rho1 or Rok knockdown in otherwise wild-type FCs (Fig. 3E,F; Fig. S1). Strikingly, however, neither Rho1 nor Rok depletion was sufficient to restore vertex opening in Tkv^QD^-expressing clones (Fig. 3J,K,M,N; Fig. S1). Likewise, expression of a dominant-negative version of the myosin II heavy chain lacking the actin-binding head domain (MHC^DN^-YFP; Dawes-Hoang et al., 2005) caused enlarged intercellular gaps in wild-type FE but failed to restore vertex opening in Tkv^QD^ clones (Fig. 3I,L,O). Thus, although TGF-β signaling induces MyoII contractility in FCs, MyoII activity is dispensable for TGF-β–mediated inhibition of vertex opening, suggesting that additional downstream effectors of TGF-β act to maintain FC vertices closed.

### TGF-**β** signaling upregulates E-Cad transcription and stabilizes it at FC vertices

We therefore investigated whether TGF-β signaling keeps vertices closed by elevating or stabilizing E-Cad at FC junctions. Before patency, E-Cad is distributed along AJs, including vertices (Fig. 4A). With the onset of patency, junctional E-Cad declines and is removed from vertices (Fig. 4B), an essential prerequisite for patency (Isasti Sanchez et al., 2021). Notably, however, E-Cad levels are elevated in anterior MBFCs, mirroring the graded TGF-β signaling activity (Fig. 4C; Row et al., 2021). This graded pattern of E-Cad levels requires TGF-β signaling, as loss of signaling in *tkv^8^* mutant clones in the anterior FE reduced junctional E-Cad levels to those observed in wild-type posterior FE, whereas *tkv^8^*clones in the posterior epithelium had no detectable effect on E-Cad levels (Fig. 4D,G). To test whether the elevated E-Cad levels in anterior MBFCs result from upregulation of *E-Cad* transcription by TGF-β signaling, we analyzed a β-galactosidase transcriptional reporter inserted into the *E-Cad* locus (*shg*-lacZ-nls; Tepass et al., 1996). Tkv^QD^-expressing cells showed 5.25-fold increased β-galactosidase signals compared to control cells (p=0.00004, n=10; Fig. 4E,H), indicating that TGF-β signaling increases *E-Cad* transcription in FCs. Consistent with this finding, depletion of the signal transducer and transcriptional effector Mad by RNAi in anterior FC clones reduced E-Cad levels to those of posterior FCs (Fig. 4F,G). Thus, E-Cad upregulation in anterior MBFCs requires canonical TGF-β signaling through Mad-dependent transcriptional regulation.

**Figure 4.**
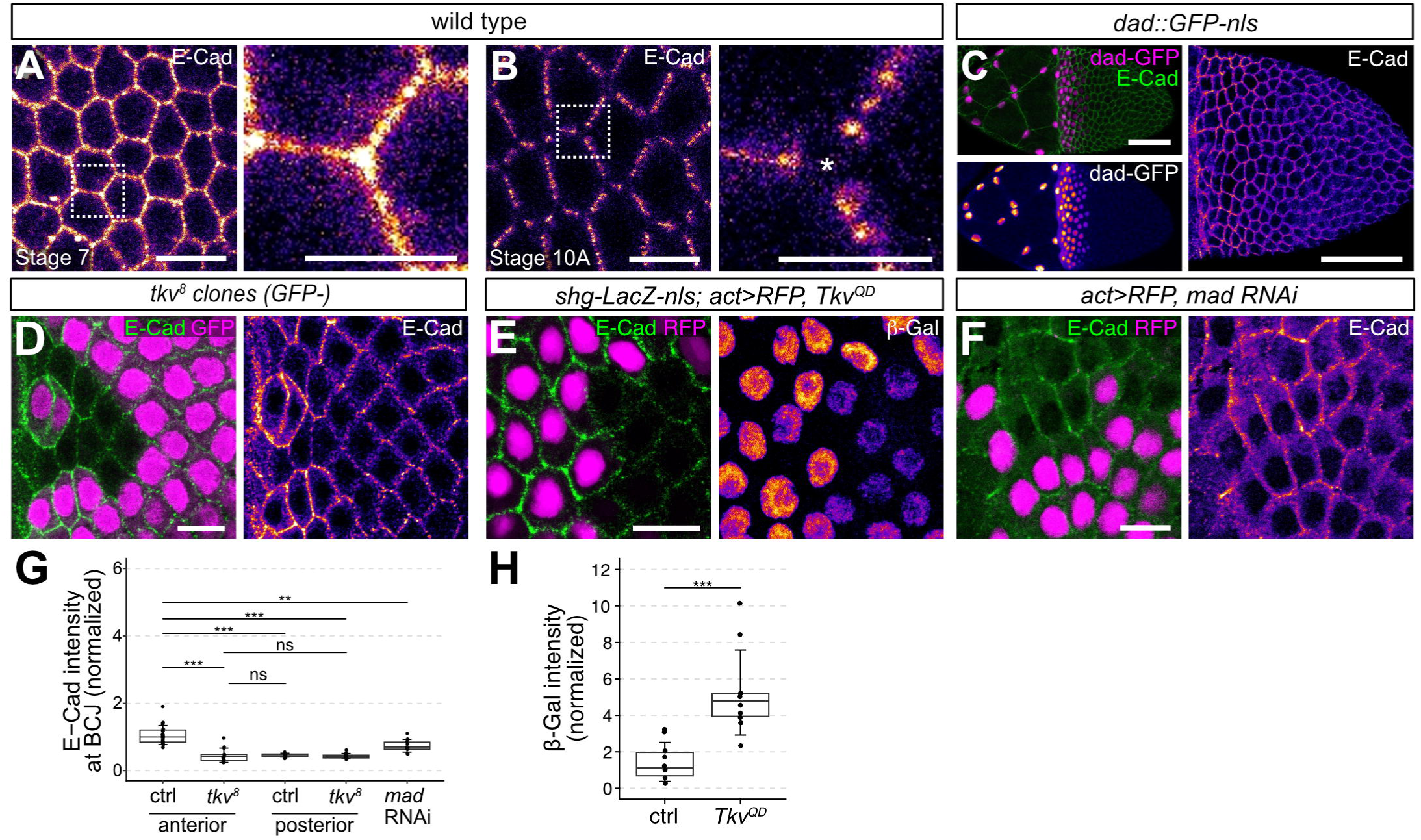
TGF-β signaling regulates E-Cadherin levels and distribution. (A-B) Basolateral sections of living pre-patent (stage 7) and patent (stage 10A) follicles expressing E-Cad::3xGFP. Note continuous junctional E-Cad signal at stage 7, whereas E-Cad is removed from vertices by stage 10A. Asterisk marks open vertex. (C) Projection of stage 10 follicle expressing *dad*::GFP-nls (magenta), stained for E-Cad (green). Single-channel images show *dad*::GFP-nls and E-Cad (intensities color-coded with heatmap). (D) Basolateral section of fixed follicle carrying *tkv^8^* clone (marked by absence of GFP (magenta)) in the anterior main body FE stained for E-Cad. Note reduced levels of E-Cad (green in merge, heatmap in single-channel image) in *tkv^8^* clone compared to neighboring control cells. (E) Basolateral section of fixed follicle expressing *shg*-LacZ-nls in all FCs and Tkv^QD^ in RFP-marked cells (magenta), stained for E-Cad (green in merge) and beta-galactosidase (single channel, heatmap). Note elevated beta-galactosidase signals in Tkv^QD^-expressing cells compared to neighboring control cells. (F) Basolateral section of fixed follicle expressing *mad* dsRNA in RFP-marked cells (magenta) in the anterior main body FE, stained for E-Cad (green, heatmap in single-channel image). Note reduced E-Cad levels in Mad-depleted cells compared to neighboring control cells. (G) Quantification of E-Cad signals (normalized) at BCJs in control and *tkv^8^*mutant cells. p-values (Pairwise Wilcoxon Rank Sum Test): ** p ≤ 0.01; *** p ≤ 0.001; ns, not significant. (H) Quantification of beta-galactosidase signals in control and Tkv^QD^-expressing cells. p-values (Welch’s two-sample t-test): *** p ≤ 0.001. Scale bars: (A-B), 10 µm, closeups 5 µm; (C), 50 µm ; (D-F), 10 µm.

Activation of TGF-β signaling in Tkv^QD^-expressing cells caused accumulation of E-Cad at BCJs (1.85-fold, p=0.0002, n=12) relative to control cells, indicating that TGF-β signaling is sufficient to upregulate E-Cad levels (Fig. 5A,I,J). In addition, Tkv^QD^-expressing FCs in stage 10A follicles retained E-Cad at vertices (TCJ:BCJ signal ratio=5.22:1; p=3.1×10^-10^; n=12), whereas E-Cad was removed from vertices of control cells (Fig. 5A,J). These effects were specific for E-Cad, as levels and distribution of Fas2 and N-Cad, which are also normally removed from FC vertices before patency (Isasti Sanchez et al., 2021), were not detectably altered in Tkv^QD^-expressing cells (Fig. S2A-D).

**Figure 5.**
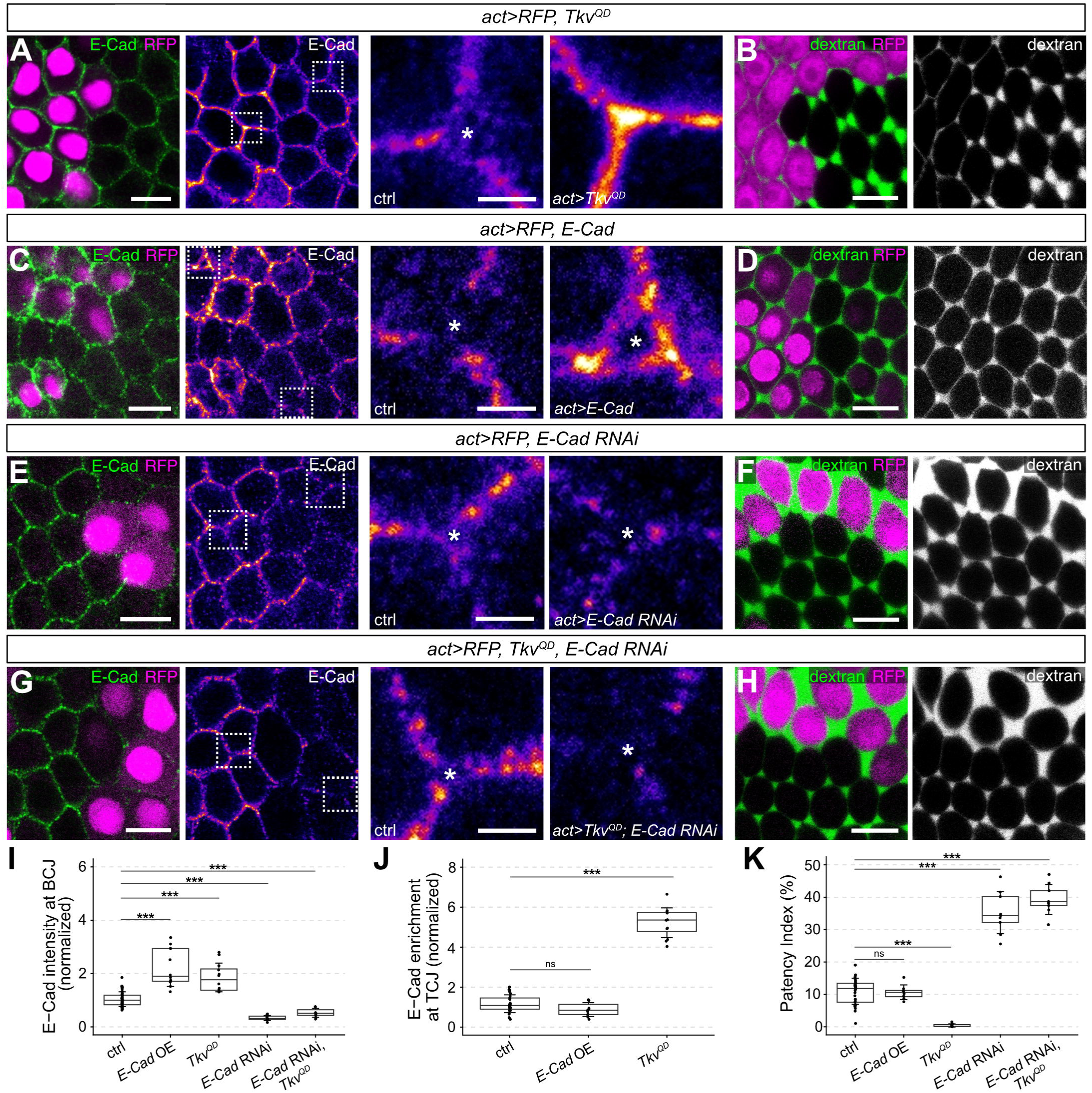
E-Cad is required for TGF-β-mediated suppression of vertex opening. (A) Basolateral section of fixed follicle expressing Tkv^QD^ in RFP-marked cells (magenta) stained for E-Cad (green; heatmap in single-channel images). Closeups show vertices of control (ctrl) and Tkv^QD^-expressing cells. Asterisk indicates open vertex after E-Cad removal. Note retention of E-Cad at vertices of Tkv^QD^-expressing cells. (B) Basolateral section of living follicle expressing Tkv^QD^ in RFP-marked cells (magenta). Dextran (green) marks intercellular gaps. Note suppression of patency in Tkv^QD^-expressing cells. (C) Basolateral section of fixed follicle overexpressing E-Cad in RFP-marked cells, stained for E-Cad. Closeups show vertices of control and E-Cad overexpressing cells. Note that E-Cad is not retained at vertices in E-Cad-overexpressing cells. (D) Basolateral section of living follicle overexpressing E-Cad in RFP-marked cells. Note that intercellular gap size is not affected by E-Cad overexpression. (E) Basolateral section of fixed follicle expressing *E-Cad* dsRNA in RFP-marked cells, stained for E-Cad. Closeups show vertices of control and *E-Cad* dsRNA expressing cells. Note reduced E-Cad levels in *E-Cad* dsRNA expressing cells. (F) Basolateral section of living follicle expressing *E-Cad* dsRNA in RFP-marked cells. Note enlarged intercellular gaps in E-Cad-depleted clone. (G) Basolateral section of fixed follicle expressing Tkv^QD^ and *E-Cad* dsRNA in RFP-marked cells, stained for E-Cad. Closeups show vertices of control cells and cells expressing Tkv^QD^ and *E-Cad* dsRNA. Note that E-Cad RNAi abolishes E-Cad vertex accumulation in Tkv^QD^-expressing cells. (H) Basolateral section of living follicle expressing Tkv^QD^ and *E-Cad* dsRNA in RFP-marked cells. Note that E-Cad depletion restores vertex opening in Tkv^QD^-expressing cells. (I,J) Quantification of E-Cad signals at BCJs (I) or TCJs (J). p-values (Pairwise Wilcoxon Rank Sum Test): *** p ≤ 0.001; ns, not significant. (K) Quantification of patency index in follicles of the indicated genotypes. p-values (Pairwise Wilcoxon Rank Sum Test): *** p ≤ 0.001; ns, not significant. Scale bars: (A-H), 10 µm; closeups 2.5 µm.

To test whether the TGF-β-induced transcriptional upregulation of E-Cad levels was sufficient to cause E-Cad retention at vertices, we overexpressed E-Cad in FCs. Strikingly, however, overexpression of E-Cad alone did not prevent its removal from vertices (Fig. 5C,I,J) and did not interfere with vertex opening during patency (Fig. 5D,K; Isasti Sanchez et al., 2021), indicating that E-Cad retention at vertices in Tkv^QD^ clones is not simply a consequence of elevated E-Cad levels. Together, these results suggest that TGF-β signaling opposes the cell contact remodeling required for patency by maintaining or stabilizing E-Cad at vertices, independent from increasing *E-Cad* expression.

### E-Cad is required for TGF-**β**-mediated suppression of vertex opening

To investigate whether TGF-β signaling modulates E-Cad dynamics to prevent patency, we depleted E-Cad by RNAi in Tkv^QD^-expressing clones. In control cells, E-Cad knockdown led to significantly enlarged intercellular gaps (Fig. 5E,F,I,K). Interestingly, E-Cad knockdown also abolished the suppression of vertex opening in Tkv^QD^ clones, yielding intercellular gaps comparable to those between otherwise wild-type FCs upon E-Cad knockdown (Fig. 5G,H,I,K). Thus, E-Cad is required for TGF-β–mediated suppression of vertex opening, consistent with a model in which TGF-β acts by preventing the removal of E-Cad from FC vertices during patency.

Similar to Tkv^QD^-expressing cells, E-Cad accumulated at FC vertices when endocytosis was blocked by expressing in FCs a dominant-negative form of clathrin heavy chain (CHC^DN^; Fig. S2E-G) or dominant-negative dynamin (Isasti Sanchez et al., 2021). We therefore hypothesized that TGF-β signaling prevents vertex opening by inhibiting the endocytosis-dependent mobility or turnover of E-Cad at AJs. To investigate the underlying mechanism, we asked whether activation of TGF-β signaling in Tkv^QD^ clones affects the distribution or levels of regulators of E-Cad dynamics. We found that the levels of p120-catenin (p120ctn), which was reported to stabilize E-Cad-based adhesion in Drosophila epithelia (Iyer et al., 2019; Myster et al., 2003), were elevated 2-fold (n=7, p=0.009) at AJs of Tkv^QD^-expressing cells, with prominent accumulation at vertices, compared to control cells (Fig. S2H,I). Overexpressing p120ctn in FC clones phenocopied the E-Cad accumulation induced by Tkv^QD^ expression (Fig. S2J,K), suggesting that p120ctn contributes to the effect of TGF-β activation in blocking patency. However, depleting p120ctn by RNAi had no detectable effect on patency (Fig. S2L,M) and did not suppress the Tkv^QD^-induced block of patency (Fig. S2N,O). Thus, p120ctn appears to contribute to, but is not essential for TGF-β–mediated inhibition of vertex opening, suggesting that p120ctn acts in a functionally redundant manner along with other factors.

### TGF-**β** signaling regulates cell-cell adhesion independently of actomyosin contractility

Finally, because cell adhesion and contractility are interlinked (Clarke and Martin, 2021), we asked whether the TGF-β-induced increase in actomyosin contractility is required to maintain E-Cad at FC vertices. We therefore analyzed E-Cad accumulation at vertices in Tkv^QD^-expressing FCs while simultaneously reducing MyoII activity by expressing dominant-negative MHC (Fig. 6). In these cells, E-Cad still accumulated at vertices (2.85-fold, p=2.84×10^-6^, n=11; Fig. 6C) despite reduced MyoII activity (pMyoII; Fig. 6A), indicating that TGF-β regulates E-Cad-based adhesion in FCs independently of actomyosin contractility.

**Figure 6.**
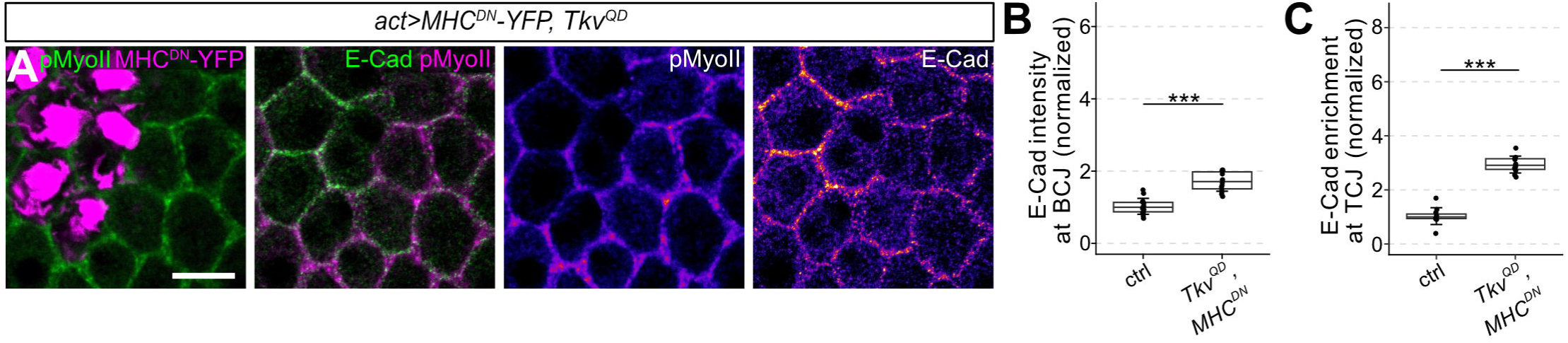
TGF-β signaling regulates cell-cell adhesion independently of actomyosin contractility. (A) Basolateral section of fixed follicle expressing Tkv^QD^ and MHC^DN^-YFP (magenta) in clones, stained for E-Cad and pMyoII. Note that MHC^DN^ expression diminishes MyoII activation (pMyoII levels) in Tkv^QD^-expressing cells, while E-Cad still accumulates at AJs and vertices. (B,C) Quantification of E-Cad signals at BCJs (B) or TCJs (C) in the indicated genotypes. p-values (Welch’s two-sample t-test): *** p ≤ 0.001. Scale bars: 10 µm.

## Discussion

Dynamic regulation of cell–cell adhesion and contractility govern epithelial plasticity during development and homeostasis, yet how upstream signals act to modulate junctional permeability remains poorly defined. Here, we dissected how TGF-β signaling regulates the adhesive and contractile machineries to control TCJ permeability during follicular patency in Drosophila ovaries. First, we show that TGF-β activity forms an anterior–posterior gradient that inversely correlates with the extent of vertex opening across the follicle epithelium. Second, TGF-β signaling suppresses patency cell-autonomously by reinforcing E-Cad–based adhesion through a twoLpronged mechanism: it induces E-Cad transcription and prevents its removal from FC vertices, in part via upregulation of p120-catenin. Third, we show that although TGF-β signaling elevates MyoII activity via the Rho–Rok pathway, the activation of actomyosin contractility is dispensable for inhibition of patency. Instead, TGF-β controls TCJ remodeling primarily by strengthening E-Cad–dependent adhesion. Thus, our findings show how an extracellular morphogen signal can tune epithelial barrier function by fortifying E-Cad-based adhesion independently of actomyosin, thus disentangling the roles of adhesion and actomyosin contractility in epithelial permeability regulation.

In Drosophila, follicular patency is initiated by downregulation of actomyosin contractility and removal of adhesion proteins from FC vertices (Isasti Sanchez et al., 2021; Jacobs et al., 2025). In the anterior FE, TGF-β signaling opposes this program by reinforcing adhesion and elevating actomyosin contractility, thereby suppressing vertex opening. These TGF-β-dependent events were shown to be essential for the cuboidal-to-squamous transition of the anterior follicle cells (Brigaud et al., 2015) and for collective tissue movements that generate the operculum and dorsal appendages at the anterior-dorsal side of the mature egg (Dobens et al., 2001; Osterfield et al., 2013). We showed that, in addition to these roles, TGF-β signaling preserves barrier integrity in the anterior FE, restricting the transepithelial passage of yolk precursors to the portion of the FE overlying the oocyte and maintaining impermeability of the FE overlying the nurse cells. This is likely to contribute to efficient uptake of yolk precursors obtained from an extraovarian source via the maternal bloodstream into the oocyte.

Actomyosin contractility and stability of AJs are generally interlinked (Clarke and Martin, 2021). Accumulation and maintenance of classical cadherins at cell junctions depends on MyoII-mediated contractility, and conversely, junctional MyoII accumulation requires E-Cad (Shewan et al., 2005; Smutny et al., 2010). Yet, surprisingly, we found that the TGF-β–induced suppression of vertex opening, albeit being dependent on E-Cad function, does not require MyoII-dependent contractility. Interfering with Rho, Rok, or MyoII activity failed to restore vertex opening in the presence of activated TGF-β signaling, whereas depleting E-Cad abrogated TGF-β–mediated suppression of patency. Thus, in this system, TGF-β signaling appears to act mainly by reinforcing E-Cad-based adhesion, independent of MyoII activity.

Our findings suggest that TGF-β signaling achieves this through at least two distinct mechanisms: (1) transcriptional upregulation of E-Cad expression, and (2) a parallel, postLtranscriptional mechanism that stabilizes E-Cad specifically at vertices and counteracts the adhesion remodeling required for patency onset. Although the exact mechanism is not yet clear, we identified the AJ-associated protein p120ctn as a factor that likely contributes to the TGF-β-induced stabilization of AJs. In mammalian cells, p120ctn stabilizes E-Cad at AJs by binding to and masking the endocytic signal in the E-Cad cytoplasmic tail (Nanes et al., 2012), thereby inhibiting E-Cad internalization and turnover (Miyashita and Ozawa, 2007). Consequently, loss of p120ctn leads to rapid E-Cad turnover (Davis et al., 2003; Xiao et al., 2005). Interestingly, interaction of p120ctn with vascular endothelial cadherin (VE-Cad) is required for normal VE-Cad levels and endothelial barrier function (Iyer et al., 2004). In Drosophila, although p120ctn is dispensable for viability and fertility, it can modulate E-Cad dynamics at AJs, possibly in a context- or tissue-specific manner (Myster et al., 2003). p120ctn was reported to modulate E-Cad endocytosis and actomyosin contractility in embryonic epithelia through opposing actions of ARF1 and Rho1 GTPase pathways (Bulgakova and Brown, 2016; Greig and Bulgakova, 2020). In the pupal wing epithelium, relocalization of p120ctn from AJs to the cytoplasm under mechanical stress was shown to promote E-Cad turnover (Iyer et al., 2019). In the FE, we showed that TGF-β signaling upregulates p120ctn and that p120ctn overexpression is sufficient to drive E-Cad accumulation at AJs; however, we found no essential requirement of p120ctn for TGF-β–mediated suppression of patency. Taken together, our findings suggest that TGF-β acts through multiple effectors, likely including p120ctn, to maintain TCJ integrity by preventing E-Cad removal from vertices. Defining the effectors that stabilize E-Cad at TCJs and their mode of regulation by TGF-β signaling are important next steps.

Cellular responses to TGF-β superfamily signals can differ widely depending on cell type, differentiation status, and developmental stage — so the same pathway can either stabilize or destabilize cell–cell junctions. In contrast to the junction-stabilizing role uncovered here in Drosophila, TGF-β signaling in mammals was reported to reduce adhesion and drive epithelial–mesenchymal transition in various contexts during development and disease (McCormack and O’Dea, 2013). In human endothelial cells, BMP6 increases barrier permeability by inducing c-Src–dependent phosphorylation and internalization of VE-Cad (Benn et al., 2016). However, even a single ligand can elicit opposite outcomes depending on cellular state: for instance, TGF-β1 has opposing effects on corneal endothelial barrier function in proliferating versus maturing cells (Beaulieu Leclerc et al., 2018). These observations underscore that ligand identity and cellular state together determine whether TGF-β signaling strengthens or weakens cell contacts.

Together, our findings identify TGF-β signaling as an upstream regulator that safeguards TCJ integrity by gating junction opening independently of actomyosin and provide a paradigm for investigating morphogen control of epithelial barrier function.

## Materials and Methods

### *Drosophila* strains and genetics

The following *Drosophila* stocks were used and are described in FlyBase, unless noted otherwise: *Act5C>CD2-Stop>Gal4 (*BDSC #4779), *hs-Flp^122^* (gift from Konrad Basler), 2x*ubi-GFP FRT40A* (Luschnig et al., 2004), *Tj-Gal4* (gift from Veit Riechmann), *dad::GFP-nls* (Ninov et al., 2010), *E-Cad::3xGFP, MyoII::3xGFP* (Pinheiro et al., 2017), His2Av-mRFP1 (Pandey et al., 2005; BDSC #23651), *tkv^8^ FRT40A (*BDSC #98353), *UAS-Tkv^QD^*(Nellen et al., 1996), *UAS-Chc^DN^ (*BDSC #26874), *shg-lacZ-nls (P{lacW}shg^k03401^*; BDSC #10377), *UAS-E-Cad-RNAi* (GD14421)v27081; VDRC #27081), *UAS-luc-RNAi* (TRiP.JF01355)attP2 (BDSC #31603), *UAS-mad-RNAi (*TRiP.JF01263)attP2 (BDSC #31315), *UAS-Rho1.RNAi.B}8.2 (*BDSC #9909), *UAS-Rok-RNAi* (TRiP.JF03225)attP2 (BDSC #28797), *UAS-MHC^DN^::YFP* (Dawes-Hoang et al., 2005), *UAS-p120ctn RNAi* (TRiP.HMC03276) attP2 (BDSC #51729).

Mosaic follicles carrying homozygous *tkv^8^* FC clones were generated using FLP/FRT-mediated mitotic recombination. Males carrying *hs-Flp^122^/Y; 2xubi-GFP FRT40A* were crossed to females carrying *tkv^8^ FRT40A/CyO*. Progeny from this cross were heat-shocked at the pupal stage (24–72 hours after puparium formation) for 3 x 60 minutes at 37°C in a water bath. For RNAi-mediated knockdown or overexpression experiments, RFP-marked flip-out clones were generated by crossing flies carrying *hs-Flp^122^* and a UAS construct to flies carrying an *act5C>CD2>GAL4* flip-out cassette and a UAS-RFP-nls transgene. Progeny from these crosses were heat-shocked as pupae (96 hours after puparium formation) for 85 minutes at 37°C in a water bath.

### Transgenic constructs

The pUAST-p120ctn construct was generated as follows. First-strand cDNA was generated from mRNA extracted from Oregon-R flies. The *p120ctn* coding sequence was amplified using forward (CAAAATGGAAGCGCGATCTCTCAA) and reverse (CCCTCTTTGCTATTGGAAT) oligonucleotides. The PCR product was cloned in the pCR-BluntII-Topo vector (Life Technologies) and excised using KpnI and XbaI. The resulting p120ctn fragment was inserted between the KpnI and XbaI sites in pUAST-attB (Bischof et al., 2007). The construct was integrated into the *attP40* and *attP2* genomic landing sites, respectively, using PhiC31-mediated site-specific integration (Bischof et al., 2007).

### Culturing follicles

Adult females were collected approximately 24 hours after eclosion and fed on yeast for two days. Flies were dissected in Shields and Sang M3++ medium (Biomol; supplemented with 1× Penicillin/Streptomycin and 10% fetal bovine serum) and mounted on glass slides with coverslip spacers. To visualize plasma membranes, follicles were mounted in M3++ medium containing CellMask Orange Plasma Membrane Stain (6.25 μg/mL final concentration; Thermo Fisher). Patency was visualized by mounting follicles in M3++ medium containing dextran (10 kDa) conjugated with either FITC or AlexaFluor 647 diluted in M3++ medium (1:200 final dilution; 25 µg/ml final concentration, Life Technologies). When required, dextran and CellMask were used together. Nuclei in living follicles were visualized using Hoechst 33342 (1 μg/ml; Sigma) diluted in M3++ medium.

### Hyperosmotic treatment

Hyperosmotic treatment was carried out as described previously (Jacobs et al., 2025). Dissected follicles were transferred to 8-well glass-bottom chambers (VWR 734-2061) coated with Poly-D-Lysine (1 mg/mL in 0.15 M H_3_BO_3_ pH 8.4). Each chamber contained 200 µL M3++ medium supplemented with dextran-AlexaFluor 647. To raise osmolarity during imaging without altering the concentration of fluorescent dextran, 10 µL 50% DMSO (final concentration 2.38% DMSO or 304 mosmol/L) were added in M3++ medium containing 37.5 µg/mL dextran-AlexaFluor 647.

### Immunohistochemistry

Female flies were collected approximately 24 hours after eclosion and were fed on yeast for two days. Ovaries were dissected in Shields and Sang M3 medium. Ovarioles were dissected and fixed using 4% PFA in 1× PBS for 15 minutes at room temperature. The ovarioles were washed three times in 0.2% PBT (PBS with 0.2% Tween 20) and then blocked with 10% BSA in 0.2% PBT three times for 30 minutes each. Samples were then incubated with primary antibodies in 10% BSA / PBT overnight at 4°C under slow agitation. After incubation with primary antibodies, samples were washed 6 times with 10% BSA for 10 minutes each at room temperature. Fluorophore-conjugated secondary antibodies (diluted 1:500 in 10% BSA) were added and incubated in the dark for three hours at room temperature. Samples were washed in 0.2% PBT six times for 10 minutes each before mounting in ProLong Gold antifade medium (ThermoFisher). To detect F-actin in fixed samples, Phalloidin conjugated with either FITC or TRITC was added along with the secondary antibodies.

For anti-phospho-Myosin staining, ovaries were dissected in Shields and Sang medium supplemented with phosphatase inhibitor solution. Dissected ovarioles were fixed in 4%PFA in 1× PBS supplemented with phosphatase inhibitor solution. The remaining steps were carried out as described above, except for the incubation with primary antibody, which was done for 24 hours at 4°C. Phosphatase inhibitor solution (modified from (Doerflinger et al., 2022)) was prepared as a 20x stock solution containing 20 mM sodium fluoride (NaF; Sigma S79209), 20 mM β-Glycerophosphate (Sigma G9422), 20 mM sodium orthovanadate (Na_3_VO_4_; Sigma 450243), and 100 mM sodium pyrophosphate (Na_4_P_2_O_7_ ⋅ 10H_2_O; Sigma S6422) and was used at 1× (1:20) final dilution.

### Antibodies

The following antibodies were used: Rat anti-E-Cad DCAD2 (1:50; DSHB), rabbit anti-phospho-Myosin Light Chain 2 (Ser19) (1:50; Cat# 3671, Cell Signaling Technology), rabbit anti-pSMAD 41D10 (1:100; Cell Signaling Technology), anti-Rho1 p1D9 (1:50; DSHB), mouse anti-N-Cadherin(1:50; DSHB), mouse anti-p120-catenin p4B2 (1:50; DSHB). Goat secondary antibodies were conjugated with Alexa 488, Alexa 568, Alexa 647, or DyLight 405 (1:500; ThermoFisher). GFP signals were enhanced with FluoTag-X4 anti-GFP (1:500; NanoTag).

### Microscopy

Imaging was performed on a Leica Stellaris Falcon confocal microscope with a 40x/1.25 NA glycerol immersion objective or on a Leica SP8 confocal microscope with a 40x/1.3 NA oil immersion objective. Image data acquired with the resonant scanner on the Leica SP8 was processed using Leica Lightning Deconvolution software (v3.5.7.23225).

### Image analysis

Images were processed using FIJI (v1.42; Schindelin et al., 2012) and OMERO (5.8.0). Image panels in figures were assembled using OMERO.figure v6.6.2 (Allan et al., 2012) and Affinity designer (v 2.6.3).

### Quantification of vertex opening

Confocal sections of the follicle epithelium overlying the oocyte were acquired at were acquired at a plane corresponding to 25% of total cell height measured from the basal surface. To quantify patency, the dextran channel image was processed with a median filter (radius = 5 pixels) and thresholded to generate a binary mask of dextran-filled intercellular gaps. The area fraction occupied by dextran signal was then calculated using the Analyze Particles function in FIJI.

### Quantification of gradients

The *dad*::GFP-nls and patency gradients were quantified by measuring nuclear *dad*::GFP-nls intensity and vertex-opening size, respectively. For the *dad*::GFP-nls gradient, fluorescence signals in MBFC nuclei and the shortest distance of each nucleus to the MBFC–nurse cell boundary were measured in FIJI. Both measurements were normalized to follicle length and plotted in R, and an exponential decay was fitted to the data. Patency index was quantified at individual vertices by drawing ROIs enclosing each dextran-filled vertex opening and calculating the dextran-filled area fraction within each ROI. The distance of each vertex from the MBFC-nurse cell boundary was measured, normalized and overlaid on the *dad*::GFP-nls-distance plot in R. The distribution was fitted with a sigmoidal function in R.

For analyzing cell shape, the dextran channel was used to segment follicle cell outlines. A “Percentile” threshold was applied to the images in FIJI and shape descriptors along with their x and y coordinates were extracted. Circularity was plotted against the x-coordinate in R and fitted with a cubic polynomial function.

### Quantification of protein levels and distribution

Protein levels at BCJs and TCJs were measured within circular ROIs (15 pixels diameter) on average intensity projections of confocal slices. Background was subtracted and intensities were normalized to the median intensity of control cells. Enrichment at TCJs was calculated as the ratio of TCJ to BCJ intensities.

### Statistics

Statistical analyses were performed in R (4.2.3) using RStudio Interface (2024.12.1+563). For phenotypic analyses, sample size (n) was not predetermined using statistical methods, but was determined by assessing the variability of a given phenotype, determined by the standard deviation. Experiments were considered independent if samples were derived from separate crosses. Each sample group was assessed for normality using the Shapiro-Wilk test and for equal variances using an F test. In case of normality and equal variances, pairwise comparisons were performed with two- or one-sided *t*-tests. In case of unequal variances, a *t*-test using pooled standard deviations was used (Welch’s two sample t-test). If data was not normally distributed, the Wilcoxon rank-sum test was applied. If samples were used for several tests, p-value were adjusted for multiple testing using the Holm step-down procedure (Holm, 1979).

## Author contributions

Conceptualization, all authors; Methodology, all authors; Investigation, H.A.; Formal Analysis, H.A.; Visualization, H.A.; Reagents and tools, H.A., T.J., M.L.; Figures, H.A.; Writing – Original Draft, H.A., S.L.; Writing – Review and Editing, S.L.; Funding Acquisition, S.L.; Supervision, S.L.

## Acknowledgements

We thank Wilko Backer for expert technical help, Markus Affolter, Konrad Basler, Yohanns Bellaiche, Christian Klämbt, Giorgos Pyrowolakis, and Ulrich Tepass for providing antibodies and fly stocks, and Ashley Hermon, Namitha Jayalal, and Archana Vellandath for comments on the manuscript. We acknowledge service and resources provided by the Münster Imaging Network, the Developmental Studies Hybridoma Bank (created by the NICHD of the NIH and maintained at the University of Iowa), and the Bloomington Drosophila Stock Center (NIH P40OD018537).

## Conflict of interest

The authors declare no competing interests.

## Funding

Work in SL’s laboratory was supported by the Deutsche Forschungsgemeinschaft (SFB 1348 “Dynamic Cellular Interfaces”), the “Cells-in-Motion” Cluster of Excellence (EXC 1003-CiM), and the University of Münster.

## Data and materials availability

Fly stocks and reagents described in this study are available upon request.

## Supplementary Material

**Supplementary Figure 1**

***Rok* RNAi diminishes Myosin II activation but does not restore patency in Tkv^QD^-expressing cells.** (A,B) Basolateral section of fixed follicle expressing *Rok* dsRNA (A) or Tkv^QD^ along with *Rok* dsRNA (B) in RFP-marked cells (magenta). Anti-pMyoII staining (green) detects active MyoII. Note reduction in active MyoII upon *Rok* RNAi both in the presence and absence of activated TGF-β signaling.

(C) Quantification of pMyoII signals at BCJs (normalized) in the indicated genotypes. p-values (Pairwise Wilcoxon Rank Sum Test): *** p ≤ 0.001.

(D,E) Basolateral sections of living follicles expressing *Rok* dsRNA (D) or Tkv^QD^ along with *Rok* dsRNA in RFP-marked cells (magenta). Dextran (green) marks intercellular gaps. Note enlarged gaps upon *Rok* RNAi both in the presence and absence of activated TGF-β signaling.

(F) Quantification of patency index in the indicated genotypes. p-values (Pairwise Wilcoxon Rank Sum Test): *** p ≤ 0.001.

Scale bars: (A, B, D, E), 10 µm.

**Supplementary Figure 2**

**TGF-**β **signaling stabilizes E-Cad at adherens junctions and upregulates p120ctn levels.** (A-B) Basolateral sections of living follicles expressing Fas2::YFP and Tkv^QD^ in RFP-marked cells (magenta). Single-channel images show Fas2::YFP. Note presence of Fas2 at vertices at stage 6 (A) in wild-type and Tkv^QD^-expressing cells, whereas Fas2 is removed from vertices at stage 8 (B) in both genotypes. Asterisks mark open vertices.

(C-D) Apical sections of fixed follicles expressing Tkv^QD^ in RFP-marked cells (magenta), stained for N-Cad. Single-channel images show N-Cad. Note presence of N-Cad at vertices at stage 6 (C) in wild-type and Tkv^QD^-expressing cells, whereas N-Cad is removed from vertices at stage 8 (D) in both genotypes.

(E-F) Basolateral sections of living wild-type follicle (E) or follicle expressing *Chc^DN^* in FCs under the control of *Tj*-Gal4. (F). Dextran (green) labels intercellular gaps, CellMask (magenta) labels plasma membranes. Note that suppression of clathrin-mediated endocytosis by *Chc^DN^* expression blocks patency.

(G) Quantification of patency index of control (ctrl) and *Chc^DN^*-expressing follicles. p-values (Welch’s two-sample t-test): *** p ≤ 0.001.

(H) Basolateral section of fixed follicle expressing Tkv^QD^ in RFP-marked cells (magenta), stained for p120ctn. Note increased p120ctn levels (intensities color-coded using heatmap in single-channel image) in Tkv^QD^-expressing cells.

(I) Quantification of p120ctn signals at BCJs in control and Tkv^QD^-expressing cells. p-values (Welch’s two-sample t-test): ** p ≤ 0.01.

(J) Apical section of follicle overexpressing *p120ctn*, stained for p120ctn and E-Cad. Note increased E-Cad levels (intensities color-coded using heatmap) at AJs in p120ctn overexpressing cells.

(K) Quantification of E-Cad levels at BCJs in control and *p120ctn* overexpressing cells. p-values (Welch’s two-sample t-test): ** p ≤ 0.01.

(L) Basolateral section of living follicle expressing *p120ctn* dsRNA in RFP-marked cells (magenta). Dextran (green) marks intercellular gaps. Note that p120ctn depletion does not affect patency.

(M) Quantification of patency index in control and *p120ctn* overexpressing follicles. p-values (Welch’s two-sample t-test): ns, not significant.

(N) Basolateral section of follicle expressing *p120ctn* dsRNA and Tkv^QD^ in RFP-positive cells (magenta). Dextran (green) marks intercellular gaps. Note that p120ctn depletion does not restore patency in Tkv^QD^-expressing cells.

(O) Quantification of patency index in control and Tkv^QD^*, p120ctn* dsRNA expressing follicles. p-values (Welch’s two-sample t-test): *** p ≤ 0.001.

Scale bars: (A-D), 10 µm, close ups 1 µm; (E, F, H, J, L, N), 10 µm.

**Supplementary Movie 1**

**Centripetal FCs resist to vertex opening induced by hypertonic treatment.** Time-lapse movie (basolateral section) of cultured stage 9 follicle expressing *dad*::GFP-nls (magenta), incubated in fluorescent dextran (green). A closeup of main-body follicle epithelium is shown. The follicle was exposed to hypertonic shock with DMSO (2.38% DMSO or 304 mosmol/L) at t=0 min. Note that addition of DMSO induces transient opening of vertices in posterior main-body FCs, but not in CPFCs expressing *dad*::GFP-nls. Time (min) is indicated. Scale bar: 10 µm.

## Notes

### Competing Interest Statement

The authors have declared no competing interest.

### Summary of Updates

Supplementary Movie S1 was added in this revised version of the manuscript.

