## Supplementary figures and images for "TGF-β signaling regulates epithelial permeability in Drosophila ovaries by modulating adhesion independent of actomyosin contractility"

### Figure S1

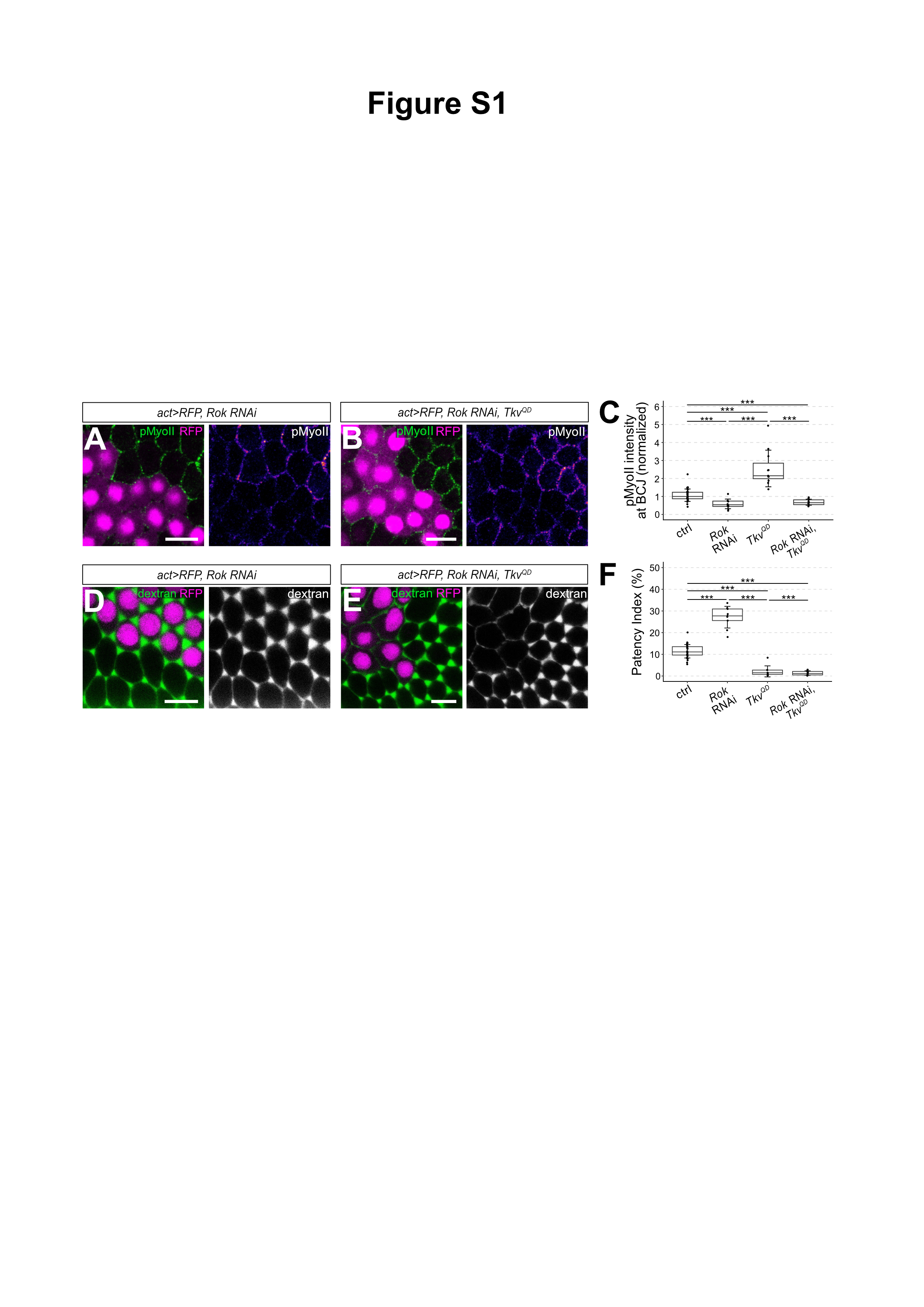

### Figure S2

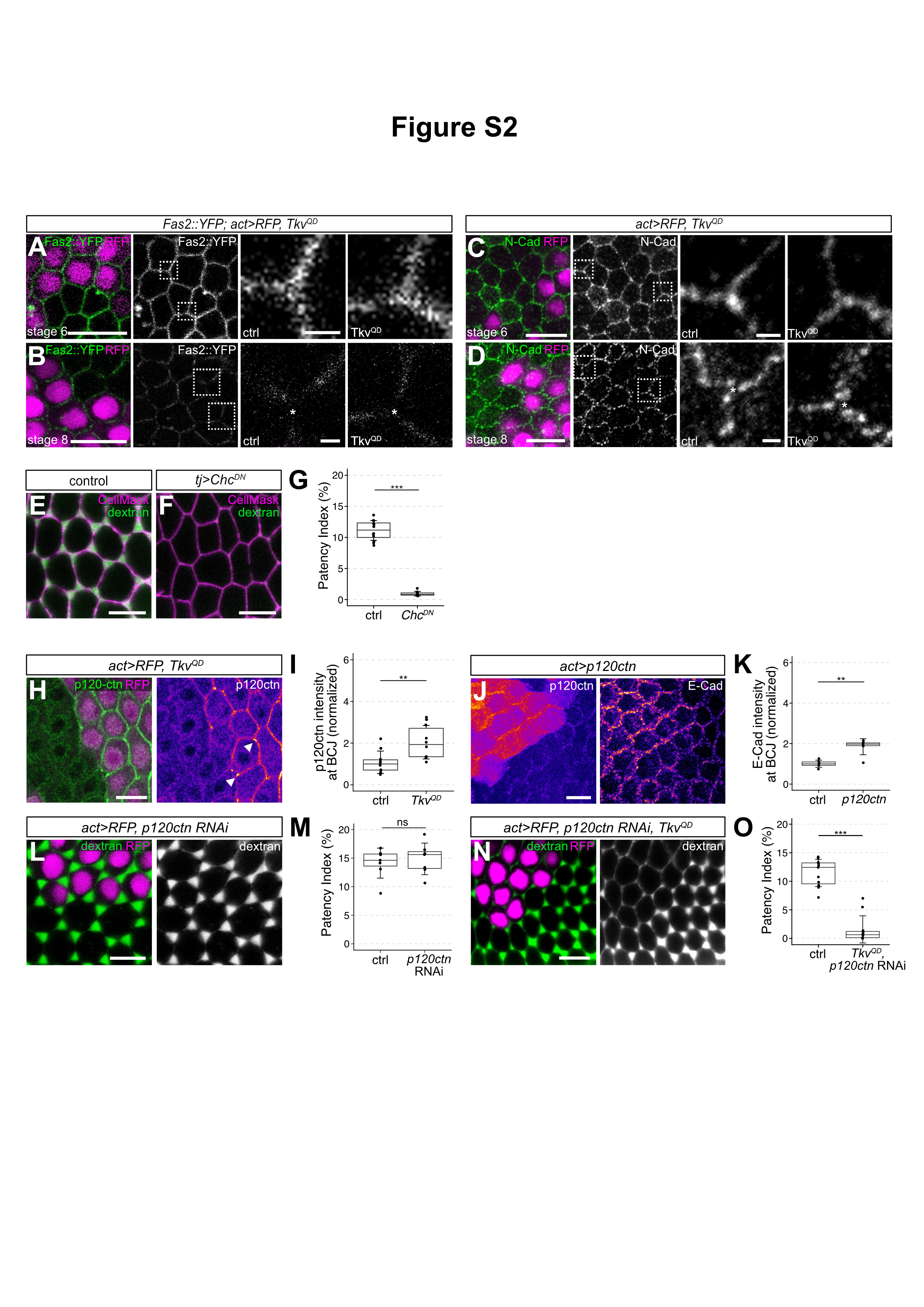
